# Pericellular oxygen dynamics in human cardiac fibroblasts and iPSC-cardiomyocytes in high-throughput plates: insights from experiments and modeling

**DOI:** 10.1101/2025.02.19.639086

**Authors:** Weizhen Li, David McLeod, Sarah Antonevich, Zhenyu Li, Emilia Entcheva

**Author notes:** **Contacts for Correspondence:** Prof. Zhenyu Li, Prof. Emilia Entcheva. W. Li and D. McLeod share first co-authorship. Note: Parts of this study were included in Weizhen Li and David McLeod’s PhD dissertations at GW.

## Abstract

Adequate oxygen supply is crucial for proper cellular function. The emergence of high-throughput (HT) expansion of human stem-cell-derived cells and HT *in vitro* cellular assays for drug testing necessitate monitoring and understanding of the oxygenation conditions, yet virtually no data exists for such settings. For metabolically active cells like cardiomyocytes, variations in oxygenation may significantly impact their maturation and function; conversely, electromechanical activity can drive oxygen demands. We used HT label-free optical measurements and computational modeling to gain insights about oxygen availability (peri-cellular oxygen dynamics) in syncytia of human induced pluripotent stem cell derived cardiomyocytes (hiPSC-CM) and human cardiac fibroblasts (cFB) grown in glass-bottom 96-well plates under static conditions. Our experimental results highlight the critical role of cell density and solution height (oxygen delivery path) in peri-cellular oxygen dynamics. The developed 3D reaction-diffusion model with Michaelis-Menten kinetics, trained on the obtained comprehensive data set, revealed that time-variant maximum oxygen consumption rate, Vmax, is needed to faithfully capture the complex peri-cellular oxygen dynamics in the excitable hiPSC-CMs, but not in the cFB. For the latter, accounting for cell proliferation was needed. Interestingly, we found both hypoxic (< 2%) and hyperoxic (> 7%) conditions can easily emerge in these standard HT plates in static culture and that peri-cellular oxygen dynamics evolves with days in culture. Our results and the developed computational model can directly be used to optimize cardiac cell growth in HT plates to achieve desired physiological conditions, important in cellular assays for cardiotoxicity, drug development, personalized medicine and heart regeneration applications.

## INTRODUCTION

Peri-cellular oxygen is the oxygen seen by the cells in their immediate extracellular space surroundings. To avoid the inherent oxygen toxicity, this peri-cellular oxygen is maintained at very low levels (physiological values in cardiac muscle vary between 3 and 6% - much lower than the 21% in ambient air or 18.5% oxygen in a standard CO_2_ incubator), and just enough to provide fuel for ATP-based energy production in the mitochondria(Place, Domann et al. 2017, Keeley and Mann 2019). Both at the cellular level and at the whole organism level, there are intricate control mechanisms in place to maintain these physiological or normoxic values as the cells have no oxygen storage on site to help respond to sudden changes in activity.

With the emergence, growth and wider use of human induced pluripotent stem cell (iPSCs) derived cells, oxygenation *in vitro* is of special interest(Sekine, Kagawa et al. 2014, Place, Domann et al. 2017, Al-Ani, Toms et al. 2018, Bagshaw, De Lange et al. 2019, Tan, Virtue et al. 2024). It was identified as a strong determinant of differentiation pathways, maturation and general functional performance. For example, with different differentiation stimuli, low oxygen tension (< 5% incubator air) promotes differentiation into either ectoderm or mesoderm lineages. In contrast, high ambient oxygen tension leads to endodermal lineages(Nit, Tyszka-Czochara et al. 2021). During differentiation of iPSC-derived cardiomyocytes (iPSC-CMs), transcription of cardiomyocytes-specific genes can be increased at lower (5%) ambient air values as compared to higher oxygen values(Ye, Qiu et al. 2020).

In post-differentiated cardiomyocytes, oxygenation is equally important for cellular metabolism and overall health, especially for metabolically active hiPSC cardiomyocytes(Ward and Gilad 2019, Ye, Qiu et al. 2020). When grown in *vitro*, in static culture, even for well-controlled incubator air oxygen, iPSC-CMs can easily transition to experiencing (peri-cellularly) hypoxia, normoxia or hyperoxia, depending on their electromechanical activity, combined with cell density and solution height. When exposed to hyperoxia, i.e. high peri-cellular oxygen levels, mitochondrial oxidative phosphorylation becomes highly active, leading to an abundance of reactive oxygen species (ROS) that can cause DNA damage. Conversely, severe hypoxia alters transcription and downregulates antioxidant defenses, leaving cells susceptible to oxidative damage and promoting increased DNA damage.

Despite its fundamental importance for cell function, peri-cellular oxygen dynamics is very rarely measured/reported. Certainly, this is not well documented or understood for the standard high-throughput (HT) glass-bottom 96 or 384 well plates widely used in stem cell research and industrial applications, such as preclinical cellular assays, cardiotoxicity testing, drug screening, growth of cells for disease modeling or regeneration.

Here we applied a high-throughput optical oxygen sensing approach(Li, McLeod et al. 2023) to study the peri-cellular oxygen changes of human cardiac fibroblasts (cFBs) and human iPSC-CMs as function of cell culture density, cell culture medium volume, and time in cell culture,. The changes in peri-oxygen levels reflect the dynamic balance of cell oxygen consumption and oxygen diffusion from the medium-air interface. We developed a 3D physics-based computational reaction-diffusion model coupled with nonlinear optimization based on the obtained comprehensive data set to match and interpret oxygen consumption behavior. Our results and the developed empirical model can directly inform future optimization of cardiac cell growth in HT plates.

## METHODS

### Human iPSC-CM and cardiac fibroblasts plating and culture conditions

Human iPSC-derived cardiomyocytes (the standard iCell Cardiomyocytes^2^ from a female Caucasian donor) from Fujifilm Cellular Dynamics International (CDI) were thawed based on the manufacturer’s instructions, then plated on top of the sterilized and fibronectin-coated oxygen sensors in 96-well glass bottom plate. Human adult ventricular cardiac fibroblasts, cFBs (Cell Applications, Inc. 307V-75a) were thawed into a T75 flask per the manufacturer’s instructions. Cells were plated into 96-well plates within five passages. Cell counting was conducted before cell plating to ensure the plating density.

The three tested cell culture densities were 156,000 cells/cm^2^, 78,000 cells/cm^2^, and 39,000 cells/cm^2^, which represented 50,000 (normal density), 25,000 (1/2 density), or 12,500 (1/4 density) cells per well (in 96-well plates), respectively, **Fig. 1A**. And four tested culture medium column heights were 9.9 mm, 6.6 mm, 3.3 mm, and 1.65 mm, which were achieved by applying 300 *μ*L, 200 *μ*L, 100 *μ*L, or 50 *μ*L culture medium per well, respectively, **Fig. 1A**. The medium exchange was done every 24 hours for hiPSC-CMs. Both hiPSC-CMs and cFBs were fixed with 3.7% formalin for immunostaining after the experiment.

**Figure 1.**
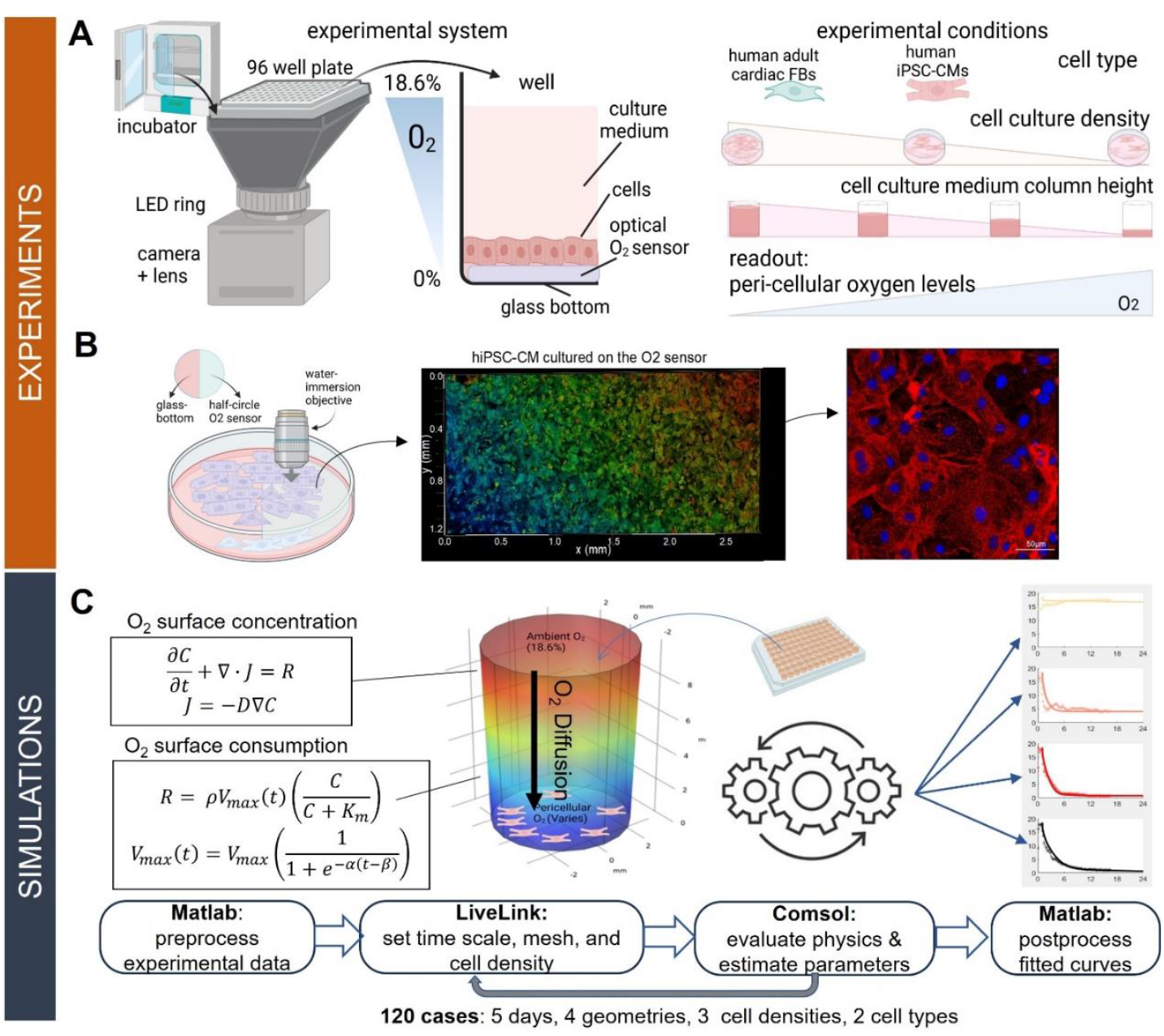
Overview of experiments and modeling of peri-cellular oxygen in HT 96-well plates. **A**. The experimental system and the experimental conditions are shown. Incubator-operated optical imaging system included a camera, a lens, LED ring and bellows extension to accommodate a glass-bottom 96-well plate. The ratiometric optical readout was calibrated in values of peri-cellular oxygen concentration over time. The experimental conditions included two cell types - adult cardiac fibroblasts, cFBs, and human iPSC-CMs, three cell densities, and four different culture medium volumes. **B**. Confirmation of cell growth on the optical oxygen sensors was done using a two-photon upright microscope with a water-immersion lens. Images of hiPSC-CMs on top of the sensor are shown at two different magnifications (25x and 40x). Color encodes depth in the lower magnification image, in which cells are labeled with Hoechst for nuclei and with ChR2-eYFP to highlight the membrane. The higher magnification image shows cytoskeleton (α-actinin) and nuclei (Hoechst); scale bar for the zoomed in version is 50µm. **C**. The computational modeling framework is a combination of 3D physics-based simulations of reaction-diffusion in the wells of a HT plate and optimization algorithms to derive empirically-based models of peri-cellular dynamics in human cFBs and iPSC-CMs, grown in glass-bottom 96-well plates, for a variety of cases. Simulations and data processing were done in Matlab, LiveLink and Comsol.

### Optical monitoring of peri-cellular oxygen in HT plates

Long-term optical monitoring of peri-cellular oxygen was done with VisiSens oxygen measuring system(Peniche Silva, Liebsch et al. 2020), as described in detail in our previous work (Li, McLeod et al. 2023), **Fig. 1A**. The optical sensors (PreSens SF-RPSu4) were laser cut into 2.5mm radius semi circles using a 30 W CO2 laser (Universal Laser Systems VLS 2.3), then mounted into a glass bottom 96-well plate (Cellvis) using vacuum tubing with their adhesive side down and sterilized with 70% ethanol over 24 hours until ethanol evaporation or removal. The sterilization process was followed by multiple PBS washes. Then the wells with the sensors were coated with 50µg/mL fibronectin overnight.

After cell plating at desired densities and solution volumes, the plate was positioned on the VisiSens system inside a humidified incubator at 37ºC and 5% CO_2_. Long-term optical measurements of pericellular oxygen were obtained by sampling the full plate in 10-min intervals, with exposure time of 0.8 s. Oxygen measurements in human cardiac fibroblasts started right after cell plating, hiPSC-CM oxygen measurements started four hours after cell plating, when cell plating media was exchanged with cell maintenance medium.

### Analysis of peri-cellular oxygen data

Peri-cellular oxygen readings were acquired as PNG images and then analyzed using the VisiSensVS software by selecting a region of interest covering the sensor in each image. The ratiometric measurements were calibrated to percentage oxygen readings using two-point calibration with 5% Na_2_SO_3_ (zero oxygen) and upon saturation with ambient air. Time-dependent calibration was applied in the first two hours of recording, considering the temperature-induced changes in transferring the plate from room temperature operation to the 37ºC humidified incubator. The data normalization was applied to the exported data to 1) equally shift the oxygen readings up to avoid negative oxygen readings; 2) apply a scale factor to the whole plate to set the maximum oxygen reading as 18.6%, the equilibrium oxygen concentration in 37ºC, 5% CO_2_ humidified incubator. Details are provided in our previous characterization study(Li, McLeod et al. 2023).

### Immunolabeling and microscopy

Human cardiac fibroblasts were labeled with Alexa Fluor™ 488 Phalloidin (Thermo Fisher A12379) and nuclei labeled with Hoechst 33342 (Thermo Fisher 62249). Human iPSC-CMs were labeled with mouse monoclonal anti-α-actinin primary antibody (Sigma-Aldrich A7811) and goat anti-mouse Alexa fluor 647 secondary antibody (Thermo Fisher A21235), Hoechst 33342 (Thermo Fisher 62249) was used for nuclei labeling. Images of human cardiac fibroblasts and human iPSC-CMs at different culture densities were taken with a Nikon Eclipse Ti2 microscope using 20x and 60x objectives. Human iPSC-CMs on top of the oxygen sensor were imaged with an upright Leica TCS SP8 microscope with 25x and 40x water immersion objective, **Fig. 1B**. Some samples were genetically modified with Channelrhodopsin-2-YFP to better visualize the cell membrane.

### Statistics and Reproducibility

Prism GraphPad was used for statistical analysis. Data are presented as individual data points with average. All experiments were repeated more than three times.

### Computational simulations of oxygen availability and consumption

A time-dependent Multiphysics reaction-diffusion model of a well in 96-well plate was built with Michaelis-Menten kinetics. The normalized peri-cellular oxygen data from human iPSC-CM and cFBs were used to feed the Michaelis-Menten kinetics (Jomezadeh Kheibary, Abolfazli Esfahani et al. 2020) for computational simulation. The COMSOL optimization module was used to define an objective function and estimate Michaelis-Menten parameters Vmax and Km. Four culture medium column heights were simulated to compare the change of Vmax under different conditions. Assumed and estimated parameters for the simulations are listed in **Table 1**.

**Table 1.**
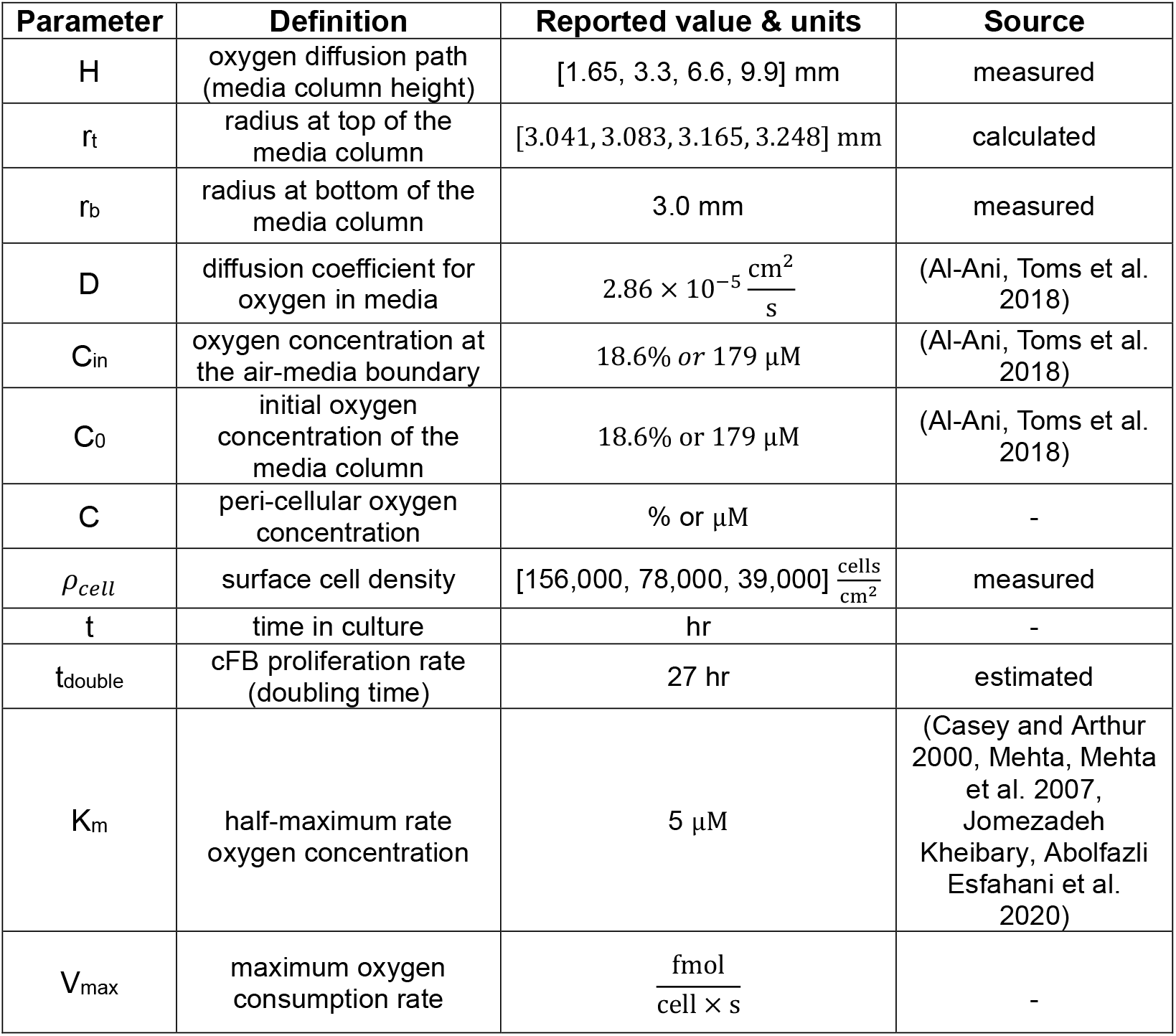
Summary of model parameters.

| Parameter | Definition | Reported value & units | Source |
| --- | --- | --- | --- |
| H | oxygen diffusion path (media column height) | [1.65, 3.3, 6.6, 9.9] mm | measured |
| $r_t$ | radius at top of the media column | [3.041, 3.083, 3.165, 3.248] mm | calculated |
| $r_b$ | radius at bottom of the media column | 3.0 mm | measured |
| D | diffusion coefficient for oxygen in media | $2.86 \times 10^{-5} \frac{\text{cm}^2}{\text{s}}$ | (Al-Ani, Toms et al. 2018) |
| $C_{in}$ | oxygen concentration at the air-media boundary | 18.6% or 179 $\mu\text{M}$ | (Al-Ani, Toms et al. 2018) |
| $C_0$ | initial oxygen concentration of the media column | 18.6% or 179 $\mu\text{M}$ | (Al-Ani, Toms et al. 2018) |
| C | peri-cellular oxygen concentration | % or $\mu\text{M}$ | - |
| $\rho_{cell}$ | surface cell density | [156,000, 78,000, 39,000] $\frac{\text{cells}}{\text{cm}^2}$ | measured |
| t | time in culture | hr | - |
| $t_{double}$ | cFB proliferation rate (doubling time) | 27 hr | estimated |
| $K_m$ | half-maximum rate oxygen concentration | 5 $\mu\text{M}$ | (Casey and Arthur 2000, Mehta, Mehta et al. 2007, Jomezadeh Kheibary, Abolfazli Esfahani et al. 2020) |
| $V_{max}$ | maximum oxygen consumption rate | $\frac{\text{fmol}}{\text{cell} \times \text{s}}$ | - |

Time-dependent simulations and parameter estimations were performed using COMSOL Multiphysics 5.5 software with LiveLink for MATLAB, and the simulation results were analyzed using MATLAB R2021b. The use of LiveLink allows for the workflow shown in **Fig 1C**. This feature allows for the scripting of COMSOL with MATLAB and automates the preprocessing and postprocessing steps required for parameter estimation and curve fitting. The 3D model used in the simulation was designed based on the dimensions of a single well in a standard 96-well plate when filled with media to a particular volume (**Fig 1C**). To study the relationship between the liquid column height and the maximum cellular consumption of oxygen, we simulated the transport of oxygen through different liquid column heights as studied. Each liquid column was initially set as fully oxygenated at 179 *μ*mol/L (or 18.5%) which corresponds to air oxygenation in a humidified 5% CO_2_ incubator at 37°C. Cell densities (*ρ*) varied between the normal, 1/2, and 1/4 densities.

The simulation was constructed to model both oxygen transport through the well and oxygen consumption at the bottom of the well. The oxygen transport equations are based on the default mass balance and molecular diffusion equations for the transport of diluted species:

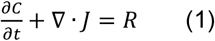

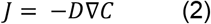

Where 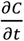 is the rate of concentration change, J is the molar flux (unit: moles per second per area) and *R* is the volume reaction rate (unit: moles per second per volume). For equation 1, the reaction term *R* is zero throughout the well volume except at the well bottom surface, where a surface reaction boundary condition is enforced (Note: for surface reactions, *R* unit is moles per second per area). The boundary condition at the well bottom surface represents oxygen consumption and was set as a modified form of the Michaelis Menten equation depending on the cell type studied. The boundary condition at the air-liquid interface was set as a constant concentration of 179umol/L for oxygen solubility in media at 37°C (Al-Ani, Toms et al. 2018). The boundary condition at the side wall was set to zero flux. Furthermore, an adjusted diffusion coefficient of oxygen in culture media at 37°C was used(Al-Ani, Toms et al. 2018).

The mesh was generated using the default normal setting with minimum element size ranging from 0.109mm to 0.178mm; minimum element quality ranged between 0.247 and 0.2841. The simulations were done on an Intel® 64bit CPU (11th Gen Intel(R) Core(TM) i7-1165G7 @ 2.80GHz, 2803 Mhz, 4 Cores, 8 Logical Processors with 16G RAM) running a Windows® 11 operating system. Cardiac fibroblast simulations were completed in 3.6 hours and cardiomyocyte simulations were completed in 25.2 hours.

### Parameter Estimation

All V_max_ parameters were estimated using a Sparse Nonlinear Optimizer (SNOPT) in COMSOL. A least-squares objective function was used, and optimality tolerance was set to 10^−9^. For cFB simulations, V_max_ was estimated as a single parameter. As a control variable, the initial value was set to 0.04 fmol/(cell*s) with lower bound 0.0005 and upper bound of 0.13 The simulated concentrations were fitted to the average sample concentrations for a given day, density, solution column height group.

For iPSC-CM simulations, time-variant V_max_ had to be implemented, and V_max_, *α*, and *β* were estimated as three parameters. V_max_ was initially set as 0.08 fmol/(cell*s) with a lower bound of 0.005 and upper bound of 0.13. The parameter *α* was initially set to 0.1 h^-1^ with a lower bound 0.056 upper bound of 0.5 while *β* was initially set to 10 h with a lower bound of 2 and upper bound of 18. The simulated concentrations were fitted to the average concentrations of samples with monotonic depletion of peri-cellular oxygen for a given day, density, column height group.

## RESULTS

The experimental measurements of peri-cellular oxygen levels were done optically with PreSens system, which contains a camera, a lens, an LED ring and an adaptor to fit 96-well plate on top(Peniche Silva, Liebsch et al. 2020, Li, McLeod et al. 2023). The LED illuminates through the glass bottom of 96-well plate and activates the oxygen sensitive dye and reference dye in the oxygen sensor. The emission lights intensity of the two dyes provides a ratio to calculate peri-cellular oxygen level based on the Stern-Volmer equation. The system can operate continuously in the incubator (**Fig. 1A**). In standard HT glass-bottom 96-well plates used in industry, as cells consume oxygen at the bottom of the well, the only source of oxygen is from the top through the solution column. An oxygen gradient establishes along the height of the well, from the air-culture medium interface at 18.6% to the cell layer at the bottom. Experimental conditions that include two types of human cells, namely human adult cardiac fibroblasts and human hiPSC-CMs, three cell culture densities and four culture medium column heights were studied to compare cell oxygen consumption dynamics.

In each 96 well, the oxygen sensor occupies half of the bottom area. The plated cells were evenly distributed across the bottom of 96-well. Half of the cells that landed on the oxygen sensor were monitored for peri-cellular oxygen changes. The other half of the cells that grew on the glass bottom enabled other bottom-up optical measurements. The cells that grew on the oxygen sensor formed well-connected syncytia, shown as membrane localized eYFP with 25x objective and cytoskeleton and nuclei structure with 40x objective in **Fig. 1B**.

A 3D reaction-diffusion model of oxygen consumption dynamics was constructed and fit to the experimental data using iterative optimization (**Fig. 1C**). The oxygen transport through the well is governed by conservation of mass and Fick’s first law of diffusion. The oxygen consumption at the bottom of the well was derived from Michaelis-Menten kinetics. The balance between oxygen diffusion through the solution column and the oxygen consumption by the cells determines the gradient of oxygen along the depth of the well as steady-state is established. Changes in culture medium height and cell density are expected to affect he dynamics of pericellular oxygen (Wagner, Venkataraman et al. 2011, Place, Domann et al. 2017, Magliaro, Mattei et al. 2019).

### Effects of cell culture density and solution height on cFBs oxygen consumption – experimental results

Cardiac fibroblasts (cFBs) are the primary cell type in the heart, providing structural support to the contracting myocardium and playing a vital role in numerous physiological signaling processes. Due to their primary function in wound healing and repair, cFBs have some/limited proliferative capacity. **Fig. 2A** shows a confluent monolayer of cFBs with the F-actin and nuclei signal in green and bule.

**Figure 2.**
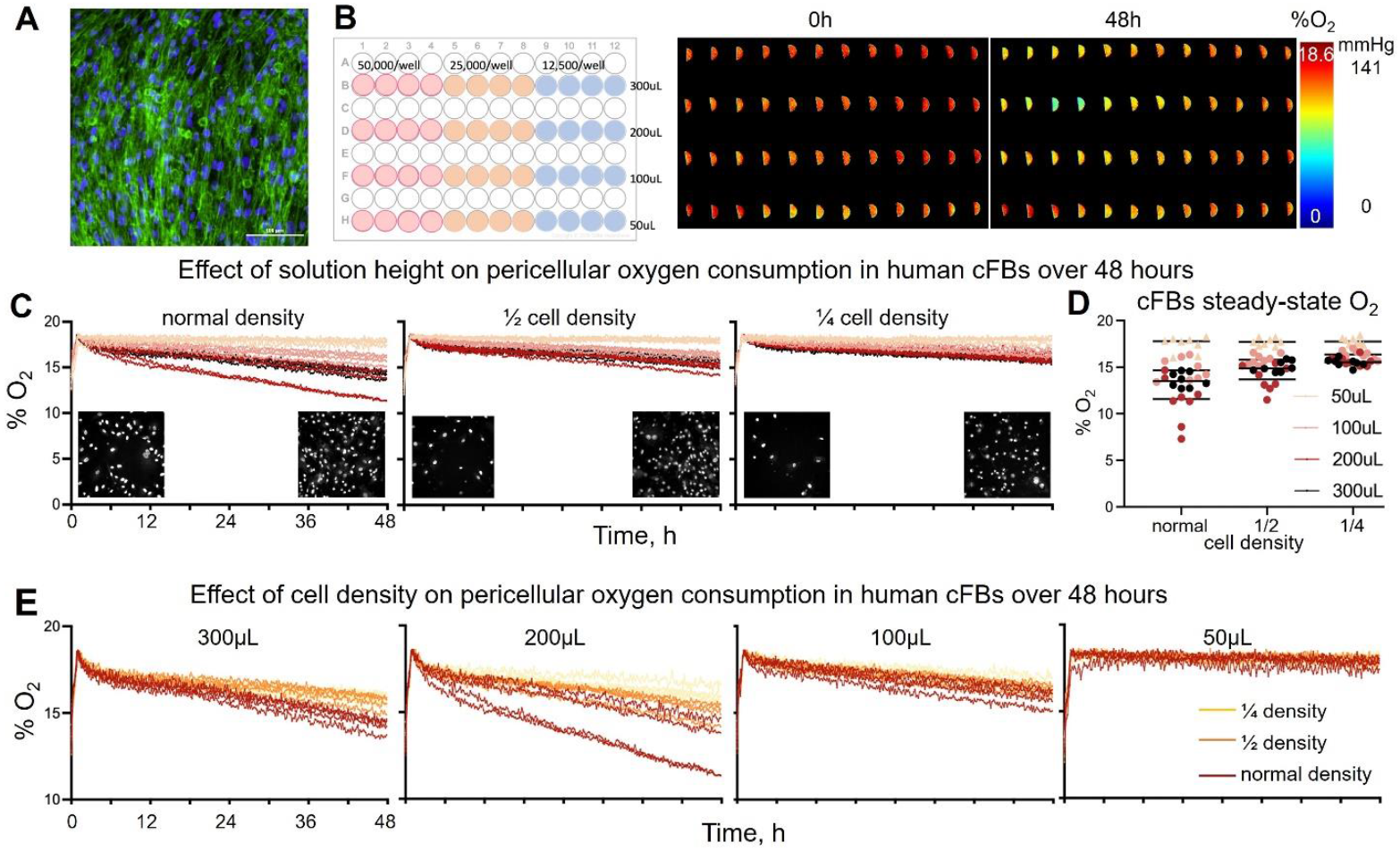
Experimental tracking of peri-cellular oxygen in human cardiac fibroblasts (cFB) over 48 hours. **A**. Fluorescence image of a confluent human cFBs monolayer (F-actin in green and nuclei in blue, with 100 µm scale bar). **B**. Experimental design with varying plating cell density (normal: 50,000 cells per well; ½ : 25,000 cells per well and ¼ : 12,500 cells per well) and oxygen diffusion path through solution volume in each well (300µL, 200µL, 100µL, and 50µL per well); four biological samples per each of the 12 conditions are included. The analyzed ratiometric images at 0h and 48h after cell plating are shown for the 48 samples. The half-circles reflect the shape of the optical sensors; color encodes % pericellular oxygen. **C**. Peri-cellular oxygen dynamics over 48 h shown for the experiment in panel B, as the samples are grouped by cell density and color encoding is used for the four solution heights. Insets are binarized images of nuclei for the different plating densities at the start of an experiment (2 h after plating) and at the 48 h endpoint. **D**. Summary plot from independent runs for steady-state peri-cellular oxygen levels for normal, ½ density, and ¼ density cFBs, n = 8 for each of the 12 cases. **E**. Peri-cellular oxygen dynamics shown for the same experiment, as the samples are grouped by solution volume/height and color encoding is used for the cell density.

In quantifying the effect of culture medium height and cell density on cell oxygen consumption, the experiment was designed as shown in **Fig. 2B**. Changes of peri-cellular oxygen levels over a 48h recording are shown in **Fig. 2B** for the studied combinations. In the 48h of cell culture, oxygen depletion was apparent in the top left quarter, for the high density (50,000 cells/ well) cell culture and high column height.

The influence of culture medium column height on cFBs oxygen consumption is shown in **Fig. 2C** under three cell plating densities over 48h. Peri-cellular oxygen of cFBs declined linearly. At the biggest column height (300 µl), there may be potential adaptation to lower oxygen levels, since the 200µl showed comparable or slightly higher oxygen depletion. **Fig. 2D** summarizes the end-point steady-state peri-cellular oxygen levels, for each condition. In 48 hours of cell culture, cFBs operated at > 10% peri-cellular oxygen levels (hyperoxic) even at the highest density (50,000cells/ well). The difference in peri-cellular oxygen decrease rate among different solution volumes remained most noticeable in the high cell density group. **Fig. 2E** shows the influence of cell density on cFB oxygen consumption, and the higher solution volume groups show most noticeable effects of cell density.

### Oxygen consumption in human cardiac fibroblasts can be modeled by time-invariant V_max_ with consideration of cell proliferation, and V_max_ is inversely dependent on cell density

For primary human cardiac fibroblasts (cFBs), the surface reaction rate was modeled using the following first order Michaelis-Menten kinetics with a proliferation term:

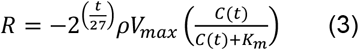

Where 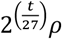 represents surface cell density as doubling every 27 hours, C(t) represents the surface concentration of oxygen at a given time point, V_max_ represents the maximum consumption rate of oxygen per cell and K_m_ represents the concentration of oxygen at half the maximum consumption rate. A value of 5*μ*M was used for K_m_ (Casey and Arthur 2000, Mehta, Mehta et al. 2007, Jomezadeh Kheibary, Abolfazli Esfahani et al. 2020), **Table 1**. Equation 3 was determined by assuming that each liquid column height would have the same doubling time. From the curve fitting, doubling time of the cFBs was estimated at 27 hours which is on par with experimentally observed. The simulated pericellular oxygen concentration was evaluated as the average concentration at the bottom boundary of the well geometry at a given time. cFB simulations were performed over 48 hours with a time step of 0.05 hours.

Without correction for doubling time, the Michaelis Menten equation did not provide good fit to experimental results as shown in **Fig 3A**. Many of the curves follow a nearly linear decline in pericellular oxygen rather than the characteristic asymptotic decline of Michaelis Menten kinetics and, therefore, a correction for cell proliferation was required for fitting. Furthermore, **Fig 3B** shows that correction using a single doubling time of 27 hours may be applied to normal, half, and quarter density samples with good results, suggesting that the correction does not depend on initial sample density.

**Figure 3.**
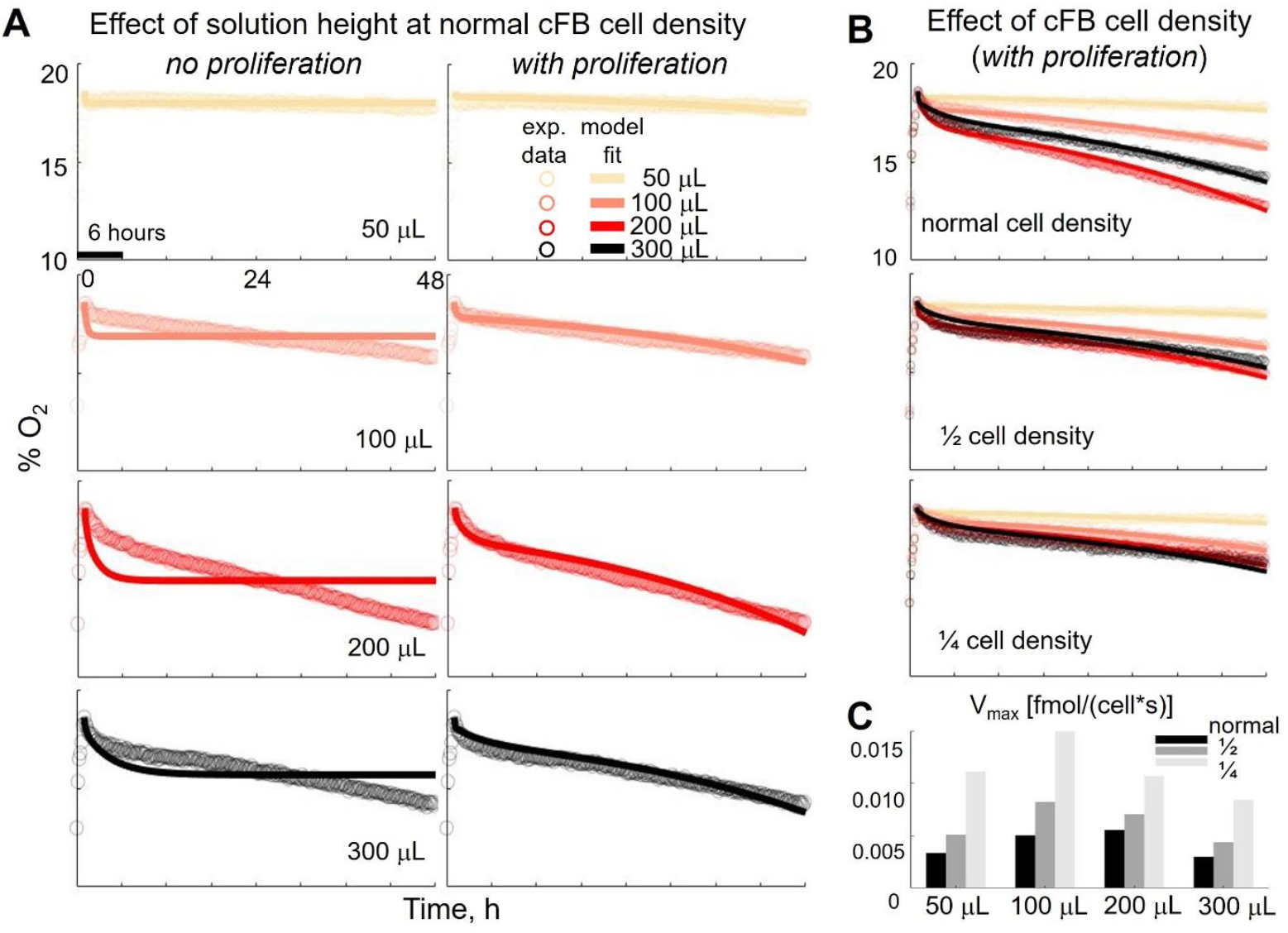
Simulations of peri-cellular oxygen in cFBs. **A**. Peri-cellular oxygen dynamics for normal cFB cell density, with the average experimental curves shown in circles and the simulation curves show in solid lines; color encodes solution height. Without proliferation in the model, fitting is poor (left). Fitting is significantly improved with a doubling time of 27 hours (right). Scale labels shown in the upper left panel apply to all graphs. **B**. Peri-cellular oxygen dynamics simulations overlaid on average experimental curves for each cell density and solution height; with proliferation, the model shows a close match with experiments across conditions. **C**. Parameter estimation for maximum cellular oxygen consumption at each density and solution column height. Estimated V_max_ (per-cell parameter) is compared across different culture medium volumes and cell densities over 48 hours of culture.

Lastly, **Fig 3C** shows that the computationally derived estimates for per-cell maximum oxygen consumption rate (V_max_, in fmol/(cell.s) appear to increase with decreasing cell density. This observation requires further investigation but may be related to cell size variations among different density cultures.

### Experimental results on the effects of cell culture density and solution height on hiPSC-CMs oxygen consumption

In **Fig. 4**, we present results on the effects of solution height and cell density on the pericellular oxygen dynamics in human iPSC-CMs over the first 24 hours after plating. Immunostaining images for α-actinin (red) and nuclei (blue) are shown under three cell culture densities (**Fig. 4A**). **Fig. 4B** shows the color-coded ratiometric images of pericellular oxygen levels at three time points (0, 12 and 24h after plating) for the experimental design/layout shown to the left with varying solution height and cell density.

**Figure 4.**
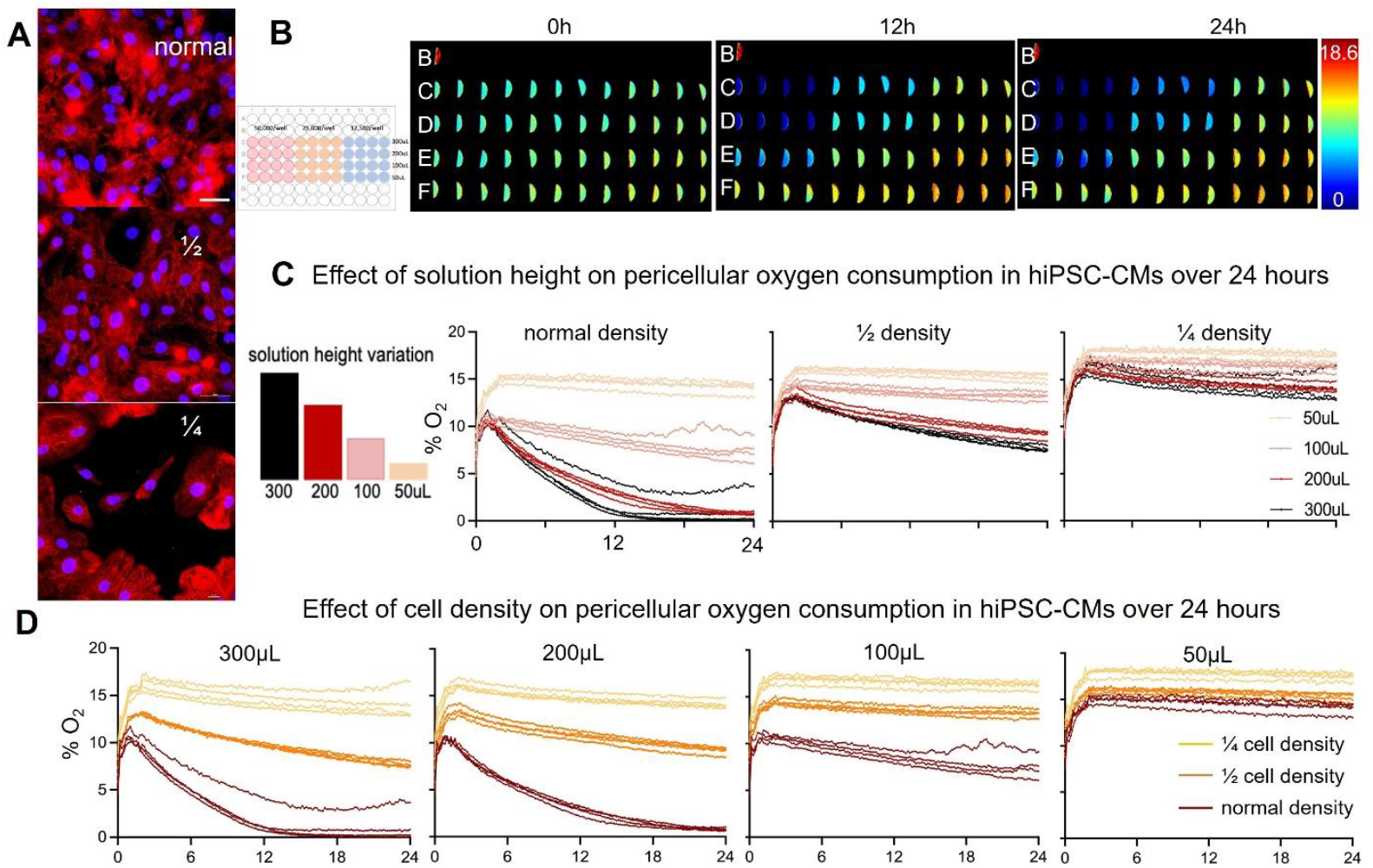
Experimental tracking of peri-cellular oxygen dynamics in human iPSC-CMs over 24 hours: influence of cell density and oxygen diffusion path (solution height) **A**. Fluorescence images of hiPSC-CMs grown at three different densities as described. Red indicates alpha-actinin and blue – nuclei. Scale bar is 50 *μ*m. **B**. The experimental design and the ratiometric images of peri-cellular oxygen for the 0, 12 and 24 hours are shown. The experimental layout and the conditions are the same as used for cFBs in Figure 2; four biological samples per each of the 12 conditions are included. **C**. Peri-cellular oxygen dynamics over 24 h shown for the experiment in panel B, as the samples are grouped by cell density and color encoding is used for the four solution heights (on the left, the solution volume – height correspondence is shown). **C**. Peri-cellular oxygen dynamics shown for the same experiment, as the samples are grouped by solution volume/height and color encoding is used for the cell density.

The measured oxygen depletion curves for hiPSC-CMs (**Figs. 4C-D**) are much more nonlinear compared to those for cFBs, with faster oxygen consumption, as expected for electromechanically active cells. And, unlike cFBs, human iPSC-CMs can easily enter a hypoxic regime (<2% pericellular oxygen) when plated at confluence in the glass-bottom 96well plates, when solution volume is 200ul or higher. Regardless of cell density, when the solution volume is low (50ul), with short diffusion pathlength for oxygen, the hiPSC-CMs appear to operate under hyperoxic conditions at >10% pericellular oxygen. As expected, both cell density and solution height (**Fig. 4C**) and cell density (**Fig. 4D**) were found to be strong modulators of pericellular oxygen levels in the human iPSC-CMs.

### Long-term adaptation of hiPSC-CM oxygen consumption – experimental results

Even differentiated human iPSC-CMs can mature further with additional days in culture. We set out to examine changes in oxygen consumption in these cells over 5 days in culture with daily exchange of culture medium to reset the oxygenation each day (**Fig. 5**). **Fig. 5A** showed the pericellular oxygen dynamics for hiPSC-CMs plated at normal cell density with four different culture medium column heights. As expected, the oxygen levels were reset to a higher level after each daily medium exchange. With advanced days in culture, the steepness of the initial oxygen depletion curve increased. The short oxygen diffusion pathlength group (50ul solution) operated at hyperoxic conditions and the balance of diffusion and consumption kept these curves relatively flat and linear. By day 5, hiPSC-CMs in the other three groups (100, 200 and 300ul) reached hypoxic levels (<2% pericellular oxygen) within a few hours after culture medium exchange. Interestingly, in addition to the monotonic depletion of oxygen (seen for cFB and for hiPSC-CMs on day 1), further days in culture revealed non-monotonic or intermittent pericellular oxygen dynamics, with partial restoration of oxygen levels after the initial depletion or with restoration and follow up depletion, **Fig. 5A**, days 2-5).

**Figure 5.**
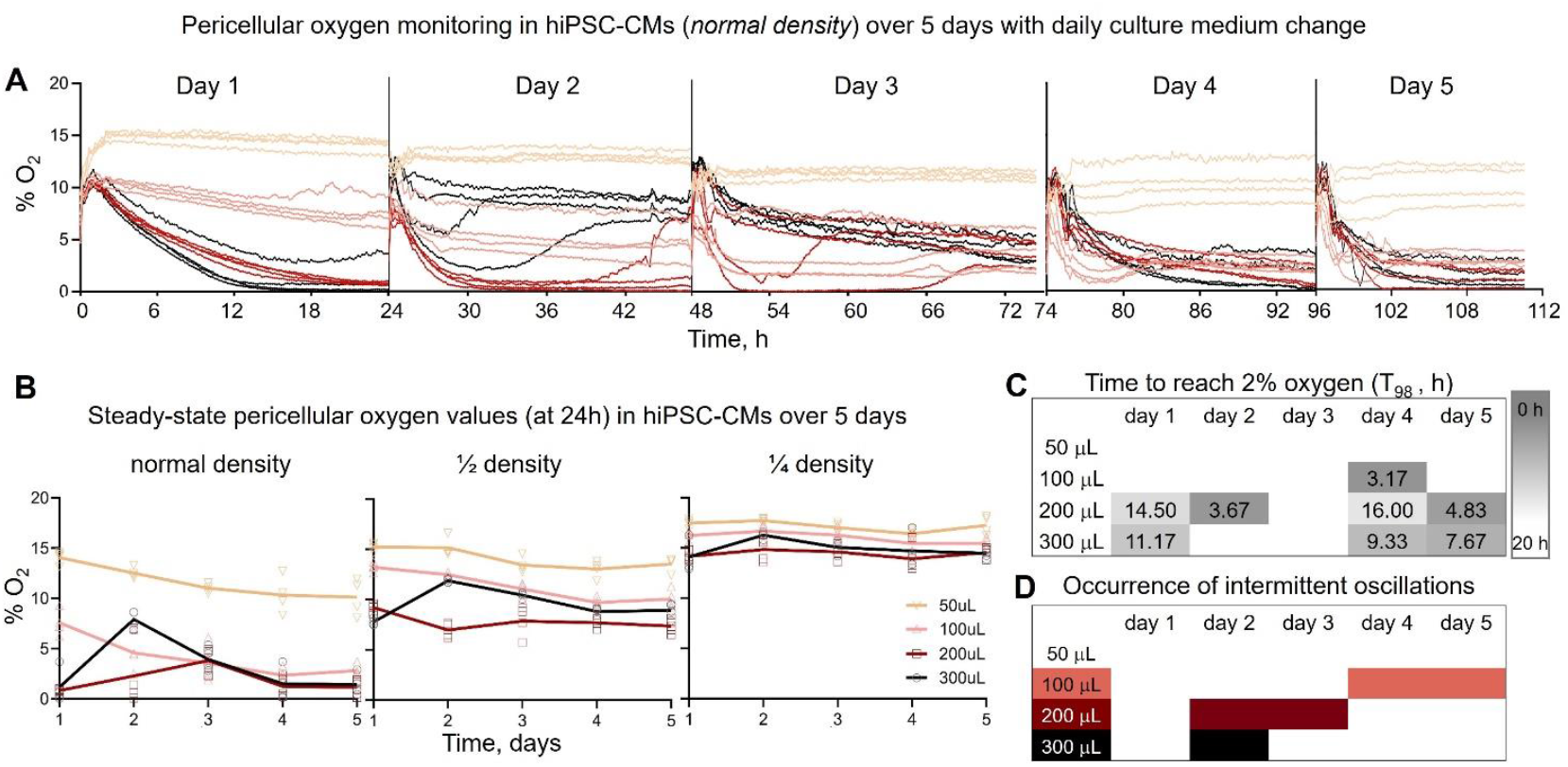
Long-term peri-cellular oxygen dynamics in hiPSC-CMs over multiple days in culture. **A**. Continuous peri-cellular oxygen reading in five days of hiPSC-CMs culture, with daily exchange of culture medium (see reset in oxygen values). **B**. The hiPSC-CMs steady-state peri-cellular oxygen over five days. “Steady-state peri-cellular oxygen” refers to the average level of oxygen measured during the last two hours of each day. **C**. Time for normal density hiPSC-CMs syncytium to reach 2% peri-cellular oxygen (T_98_, h) across five days of cell culture. **D**. Occurrence of intermittent oscillations of oxygen consumption over five days of hiPSC-CM culture.

The progressively faster initial oxygen depletion and lower steady-state values (summarized in **Fig. 5B**) likely indicate cell maturation with extended cell culture duration. For the sparser plating, esp. for the ¼ cell density group, the overall oxygen consumption rate was low and, regardless of solution volume, the steady state pericellular oxygen remained at hyperoxic levels, similar to cFBs. The initial oxygen depletion rate for all studied combinations is summarized in **Fig. 5C** as time to reach 2% (average values for the group). In some cases, hiPSC-CMs reach hypoxic values within 3-4 hours after media exchange, particularly for the intermediate solution volumes of 100ul and 200ul. As expected, the low solution volume of 50ul kept the pericellular oxygen levels high and never reaching hypoxic values.

**Fig. 5D** summarizes the occurrence of intermittent oscillations in pericellular oxygen, starting on day 2 for the high solution volumes, which would be more likely to lead to hypoxic conditions (300ul and 200ul), but also for 100ul on days 4 and 5. These oscillations are not linked to disturbance in the solution or readings, since they are measured in select wells while the rest of the samples do not show them. We first reported on this observation in human iPSC-CMs (but not human cFB) in an earlier report(Li, McLeod et al. 2023). Here we show that this is a dynamic process of likely adaptation that evolves differently over time depending on solution levels.

### Modeling oxygen consumption in human iPSC-CMs – time-variant Vmax

In modeling the oxygen consumption of human iPSC-CMs, we focused here only on the monotonic depletion curves, since the intermittent dynamics is a new observation that requires further experimental data and mechanistic analysis. As human iPSC-CMs do not proliferate, the cell density was set constant. For a good fit of the experimental data, it was necessary to use a modified time-dependent V_max_:

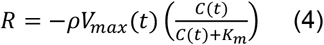

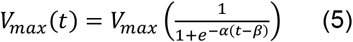

Where *α* and *β* are parameters used to describe a time-dependent maximum oxygen consumption rate. A value of 5*μ*M was used for K_m_ (Casey and Arthur 2000, Mehta, Mehta et al. 2007, Jomezadeh Kheibary, Abolfazli Esfahani et al. 2020), **Table 1**. The simulated pericellular oxygen concentration was evaluated as the average concentration at the bottom boundary of the well geometry at a given time. iPSC-CM simulations were performed over 24 hours with a timestep of 0.05 hours.

The form of equation 5 was determined by assuming that, for a given day, V_max_ could increase before saturating to a given value. In trying to fit the experimental peri-cellular oxygen data with a Michaelis-Menten dynamics, we observed that for all column heights on the first day, fitting was poor with the standard Michaelis-Menten equation. For each column height, the time to reach steady state was longer than would be expected for following the unmodified MM equation. This poor fit was also observed for many column heights over the remaining days. By assuming that the cell density remains constant, increasing the Vmax term within the model allows for improved fitting, **Fig. 6**. We note that V_max_ cannot continue to increase indefinitely; a sigmoidal curve is often used to capture physical/biological processes. Therefore, the sigmoidal curve shape was chosen to capture the varying degrees of increasing V_max_ for different density, days, column height conditions while allowing for saturation.

**Figure 6.**
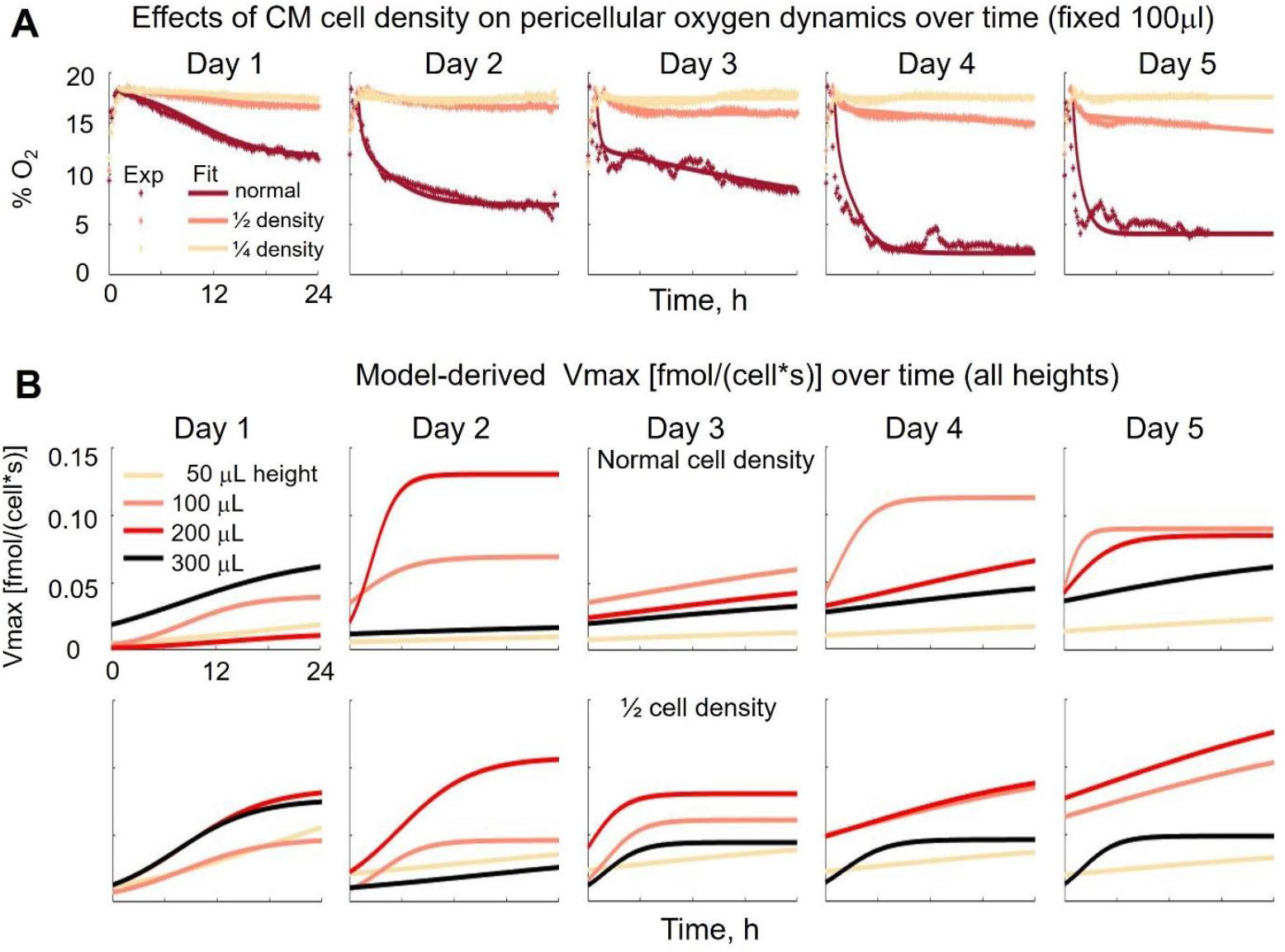
Simulations of peri-cellular oxygen dynamics in hiPSC-CMs over multiple days in culture and derivation of time-variant V_max_. **A**. Simulated peri-cellular oxygen concentrations for 100 µL media volume across cell densities and days in culture. By accounting for the consumption dynamics (through Vmax), good fitting can be achieved for each density over five days. Note the progressively faster initial depletion of oxygen. **B**. The time-variant V_max_ values from empirically based optimization are plotted for normal and half density samples across conditions; the low-density samples were well described by time-invariant V_max_, similar to the cFBs. V_max_(t) varied with column height and day. Axes shown for the upper left plot in each panel apply across plots for that panel.

From **Fig. 6B**, the smallest column height (50uL) exhibits minimal change in V_max_ over time to achieve a good fit. This condition (of high oxygen availability) likely can be modeled with minimal changes in V_max_. However, other conditions, particularly on day one, exhibit a steady increase with a plateau region and this behavior is more apparent in samples at half density. This could be due to the rapid depletion of oxygen in samples at normal density and higher column heights (200uL and higher). Particularly for the 300uL, the normal density case does not exhibit much change in V_max_ over time after day 1, as the cells may be adapting to the hypoxic conditions. The intermediate volume cases of 100uL and 200uL display the most complex dynamics, as they have greater oxygen availability without being over oxygenated and therefore may exhibit more variability in oxygen consumption to maintain healthy metabolism. Further investigation is required to elucidate the mechanisms behind the time variation in oxygen cellular oxygen consumption.

### Summary of results and analytical solution for steady-state pericellular oxygen in static culture in glass-bottom HT plates

In **Fig, 7**, we summarize the findings from this study as factors affecting the balance between oxygen diffusion and oxygen consumption by the cells. By assuming that the pericelluar oxygen concentration can reach a steady state (i.e. dC/dt = 0), and using two additional approximations: 1) the well has a uniform diameter; 2) the cell monolayer can be approximated as an O_2_ consuming surface with a uniform consumption rate *R*, we can derive an analytical solution of the pericellular oxygen concentration, governed by the following equation (the detailed derivation is given in the **Supplement**):

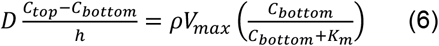

Equation 6 can be solved for pericellular oxygen C_bottom_ using the quadratic formula and keeping the positive root. Equation 6 relates the supply of oxygen through diffusion to the demand of oxygen from cellular consumption. Supply from diffusion is 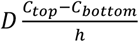 while demand from cellular oxygen consumption is 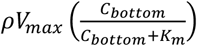. The interplay between these two phenomena is displayed graphically in **Fig. 7A**. The balance of supply and demand can be adjusted by altering certain parameters such as liquid column height and cell density. Eq. (6) may also be used to sweep parameters such as V_max_ and cell density as pericellular oxygen changes. **Fig. 7B** shows the results of one such sweep for V_max_ where the supply curve (blue) is plotted as the left side of (6) and the demand curve (red) is plotted as the right side of (6). We have highlighted regions of hypoxia, normoxia and hyperoxia that can be entered by various combinations of solution height and cell activity (for a fixed confluent cell density).

**Figure 7.**
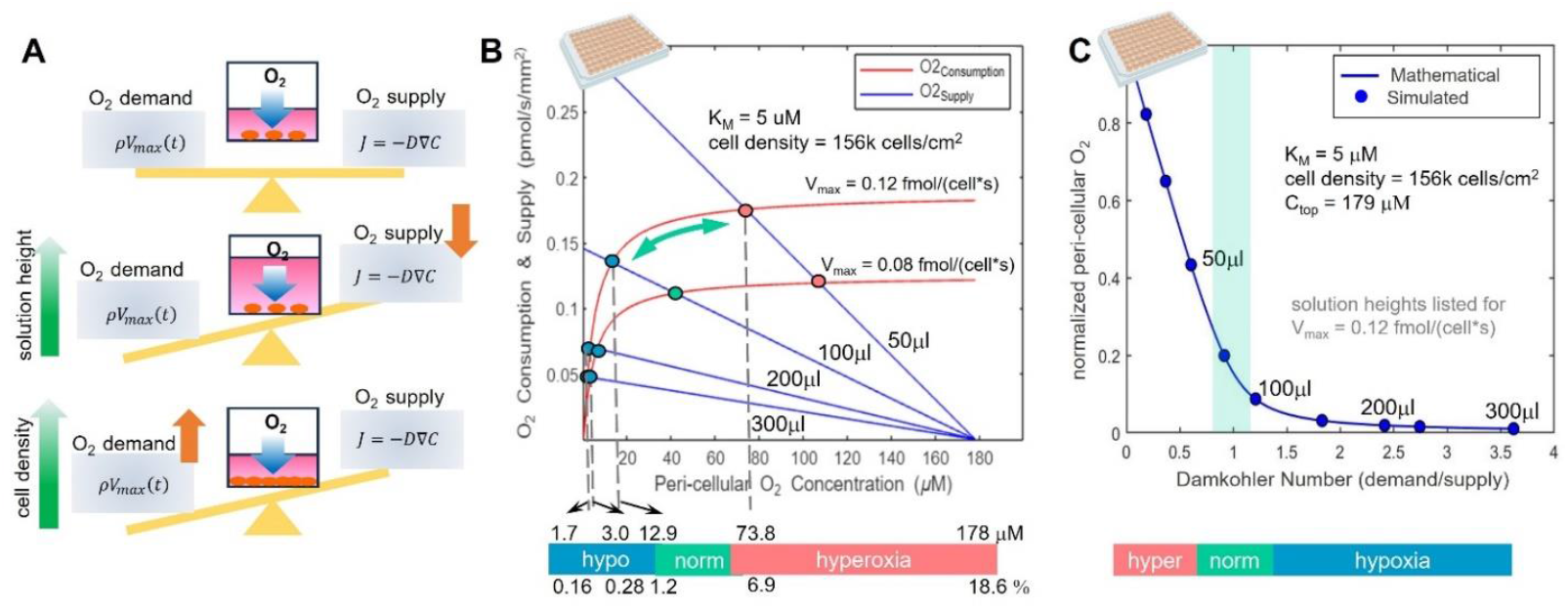
Summary illustration of peri-cellular oxygen dynamics in human cardiac cells. **A**. Schematic of how cell density and medium column height influence the balance of peri-cellular oxygen supply and demand. Increasing solution height or cell density decreases the supply or increases the demand, respectively, leading to lower levels of pericellular oxygen at steady-state. **B**. peri-cellular oxygen consumption (red) and supply (blue) curve of normal density hiPSC-CMs under four different medium column heights. Depending on Vmax, the crosspoints indicate the peri-cellular oxygen steady-states for each condition; these are color-coded to indicate predominant operation in hyperoxic, normoxic or hypoxic conditions. **C**. Normalized steady-state pericellular oxygen simulated (dots) and plotted from a dimensionless analytical solution (line). The system pericellular oxygen may be understood from the Damkohler number where values near 1 indicate normoxic conditions and parameters such as solution volume, cell density and ambient oxygen concentration may be adjusted to achieve a desired steady-state regime. For the fixed parameters for hiPSC-CMs shown, such normoxic conditions are seen for media volumes between 75 and 100 µL.

Equation 6 can also be presented in a dimensionless form, by using the Damkohler number *Da* (defined in equation 8 below) which is the ratio of reaction rate versus diffusion rate)(Truskey, Yuan et al. 2009). This dimensionless form of eq. (6) can be obtained by dividing eq. (6) by 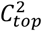 and applying the following dimensionless relationships:

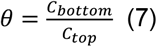

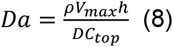

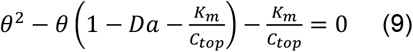

This equation was solved using the quadratic formula, like (6), while sweeping along varying Damkohler number values to generate the curve shown in **Fig 7C**. The Damkohler numbers shown were varied only according to liquid column height while V_max_ was kept constant. From the curve shown in **Fig 7C**, we can see that for Da << 1 or Da >> 1 the samples will be hypoxic or hyperoxic, respectively. Da can be estimated using cell type, metabolic state, cell density, incubator oxygen concentration, and liquid column height, and can be used as a simple indicator of oxygenation status. The analytical solution shows nearly exact agreement with simulations for the steady-state case. Varying V_max_, ρ, the diffusion coefficient, or liquid column height does not change the shape of the curve shown in **Fig. 7C**. Therefore, the Damkohler number may be used to compactly describe many tunable system parameters with a single number that offers a quick insight into the steady-state pericellular oxygen regime (hypoxic, normoxic, or hyperoxic).

## DISCUSSION

Human iPSC-CMs are a valuable cardiac experimental model due to their scalability and relevance to human physiology. Even though high-throughput formats, like 96-well and 384-well plates, are widely used in industry for hiPSC-CMs-based studies cardiotoxicity assessment, genetic modifications, disease modeling, expansion for regenerative applications etc., the oxygenation state of samples in these formats has not been fully characterized. With recent (2022) regulatory changes by the FDA for modernization (Ahmed, Shivnaraine et al. 2023), the importance of such human *in vitro* HT cellular assays will only be increasing in preclinical testing for drug development. Therefore, understanding of the oxygenation conditions is important for achieving more physiological results.

Unlike in a polystyrene petri dish, in the HT glass-bottom well plates, the medium-air interface is further away from the cells and the top surface serves as the only source of oxygen for the cells because of the oxygen non-permeable bottom. For static culture, which is the norm, simple diffusion through the solution column may or may not balance the oxygen demands of the cells, depending on their plating density, the solution column height and the cell activity (Sekine, Kagawa et al. 2014, Peniche Silva, Liebsch et al. 2020, Veldhuizen, Chavan et al. 2022). Recent optical methods of directly assessing pericellular oxygen dynamics continuously and label-free(Tschiersch, Liebsch et al. 2012, Rolletschek and Liebsch 2017, Peniche Silva, Liebsch et al. 2020, Grün, Pfeifer et al. 2023, Li, McLeod et al. 2023) allow to approach this problem in a high-throughput format. Careful calibration can yield insights on conditions of hypoxia, normoxia or hyperoxia as defined for the peri-cellular space and not for the bulk dissolved oxygen(Place, Domann et al. 2017, Keeley and Mann 2019).

Here, we focused on two types of human cardiac cells – the non-excitable cFB and electromechanically active hiPSC-CMs. Data on pericellular oxygen dynamics in these cells grown in HT plates is extremely scarce, practically non-existent(Li, McLeod et al. 2023).Therefore, we set out to obtain a rich experimental data set using optical means in HT 96-well plates and developed 3D computational models to fit and interpret the data.

There were some expected results, e.g. increased cell density and increased oxygen diffusion pathlength (solution volume) led to more likely hypoxic conditions (likelihood for pericellular oxygen to drop to <2%) in the electromechanically active hiPSC-CMs, **Figs. 4-6**, but not in the adult human cFBs. In fact, for all studied conditions, the cFBs were at what would be considered hyperoxic conditions (>10% pericellular oxygen), **Figs. 2-3**. In this study, we did not specifically modulate their transition to myofibroblasts (a more contractile phenotype)– neither accelerated or countered such transition(Kostecki, Shi et al. 2021, Song, Alvarez-Laviada et al. 2023). However, it would be interesting to apply the developed assays and models to see if noninvasive monitoring of peri-cellular oxygen dynamics can effectively distinguish different states of human cardiac fibroblasts, considering their key role in cardiac disease progression(Kohl and Gourdie 2014, Rog-Zielinska, Norris et al. 2016).

An extreme shortening of the diffusion pathlength (50ul of solution per well), which is close to conditions experienced in standard polystyrene dishes with oxygen-permeable bottom, resulted in what would be considered hyperoxic conditions not only for cFBs but also for human iPSC-CMs regardless of cell density. This phenomenon – the likelihood to have hyperoxic conditions even for very energetically-demanding cells – is little appreciated(Keeley and Mann 2019). Hyperoxia is in some respects more detrimental to the cells compared to hypoxia, considering that they do not encounter it within the body and have no well-developed defenses.

At the other end of the spectrum, for electromechanically active human iPSC-CMs, we report on easily reachable hypoxic conditions, even shortly (3-4 hours) after culture media exchange, under normal solution levels, such as 200ul or higher, **Figs. 5-6**. We surmise that under active electrical or mechanical stimulation or upon adrenergic stimulation or other inotropic activation, hypoxic conditions will be reached even quicker inevitably in such static cultures in HT plates. This is something that needs to be considered in drug testing or other characterization studies, because the altered metabolism can impact the readouts and may not reflect *in vivo* conditions, where blood flow regulation insures a feedback control of physiological pericellular oxygen levels.

Overall, cell density and solution height were confirmed as critical modulators of pericellular oxygen dynamics in both cardiac cell types studied here. Furthermore, several insights were reached when a 3D reaction-diffusion model was constructed and fit to the rich empirical dataset. Specifically, the model may be used as an indirect online estimator of cell proliferation for cFBs, as V_max_ in the Michaelis-Menten kinetics needed to be modified with such cell proliferation term, and that term can be extracted upon a good fit of the oxygen consumption curves. The sparser cFB cultures had higher per-cell V_max_, likely because of larger cell volume and/or higher metabolism for these cells when they have room to spread, **Fig. 3C**. More interestingly, we found that time-variant V_max_ was a necessary part of the computational model to successfully match the highly nonlinear profiles of oxygen depletion by the hiPSC-CMs, **Fig. 6B**. This may reflect adaptation to culture medium exchange, and to oxygen availability based on solution height. In general, the smaller solution volume samples (50ul), which likely created hyperoxic conditions, had the lowest V_max._ These curves were followed by the highest solution volume group, 300ul, which likely created hypoxic conditions due to the long oxygen diffusion pathlength. Interestingly, intermediate 100 – 200ul solution volumes yielded the highest V_max_ curves – likely an indication of optimal metabolic conditions for the human iPSC-CMs.

Long-term observations (over five days) showed that hiPSC-CMs exhibited progressively steeper initial slope of oxygen depletion – a likely sign of maturation with time in culture, **Fig. 5**. This higher activity also led to more easily achievable hypoxic conditions (<2% pericellular oxygen) even for 100ul solution volume. The tools developed in this study can be deployed and validated for monitoring the differentiation and maturation of human iPSC-CMs. Model-based interpretation of parameters can yield a metabolic assessment tool that is simpler and more scalable compared to NADH-based monitoring of metabolism, for example(Desa, Qian et al. 2023). Furthermore, for this static culture, the measured pericellular oxygen values can easily be converted into oxygen consumption rate (*OCR*), more commonly reported in the terminal Seahorse assays(Gerencser, Neilson et al. 2009). This can be done by considering the solution height and the oxygen gradient.

One of the more interesting new results from this study is the non-monotonic intermittent dynamics for pericellular oxygen in the hiPSC-CMs after day 2 in culture, **Fig. 5A, C**, first reported by us earlier(Li, McLeod et al. 2023). After the initial rapid oxygen depletion, in some cases, the oxygen levels were partially restored or exhibited slow oscillations, likely indicative of slowing down of electromechanical activity through some adaptation process to counter the hypoxic conditions. We consider this to be a likely explanation because electromechanical activity accounts for >80% of the oxygen consumption in cardiomyocytes(Keeley and Mann 2019), and because there was no solution disturbance in the plate. The general lack of prior pericellular oxygen level data over time in culture from cardiomyocytes or other excitable cells, leaves us with a desire to probe these dynamics more closely in future investigations. A limitation of this study is that our computational modeling only dealt with the monotonic curves of oxygen depletion due to the need to acquire more data and establish the factors that promote non-monotonic responses. We believe that such line of inquiry can yield valuable insights on the adaptation mechanisms in cardiomyocytes to hypoxic conditions, and specifically how human iPSC-CMs adjust their activity when encountering such conditions. This is particularly relevant to transplantation for heart repair in the early stages of integration, before blood supply to the transplants is established. This is also the time period in which most transplantation triggered arrhythmias occur, which may have relation to the oxygenation/metabolic conditions(Barker, Carpenter et al. 2022, Marchiano, Nakamura et al. 2023).

The presented experimental approach here offers a label-free longitudinal method for assessment of oxygenation and metabolism in cardiac cells grown in HT plates. We have adopted the method(Li, McLeod et al. 2023) to make it compatible with other electromechanical functional measurements done by all-optical electrophysiology (Klimas, Ortiz et al. 2020, Heinson, Han et al. 2023), and for follow-up molecular quantification in the same samples(Li, Han et al. 2022, Han, Heinson et al. 2023, Liu, Han et al. 2023). This provides a powerful pipeline for characterization of human cardiac cells. Further developments can include addition of in-plate perfusion(Wei, Li et al. 2020) and oxygenation, as needed, based on model prediction. As a practical illustration of our key insights to inform optimal growth of cardiac cells in HT plates, we provided a simple analytical solution and graphical illustration in the **Supplement** and in **Fig. 7**, for the steady-state in static culture. Oxygenation and metabolic conditions are known modulators of electromechanical activity and such insights can augment new approaches to cardiotoxicity testing and drug development.

## ACKNOWLEDGEMENTS

This study was supported in part by grants from the NIH-NHLBI R01HL144157 (to EE and ZL) and from the NSF EFMA1830941 (to EE and ZL).

## AUTHOR CONTRIBUTIONS

WL and DM contributed equally to this work. WL performed most experiments, analyzed the data and produced the figures. SA performed experiments with cardiac fibroblasts and analyzed the data. DM and ZL built the computational model, performed the simulations and solved the analytical equations. EE and ZL designed and oversaw the study and secured funding. WL and EE wrote an initial draft, which was edited with input from DM, ZL and SA.

## DATA AVAILABILITY

All data are included in the manuscript.

## SUPPLEMENTARY INFORMATION

### Derivation of the analytical solution of steady-state pericellular oxygen in static well culture

In the following derivation, we assume that the pericelluar oxygen concentration can reach a steady state (i.e. dC/dt = 0), and two additional approximations: 1) the well has a uniform diameter; 2) the cell monolayer can be approximated as an O_2_ consuming surface with a uniform consumption rate R.

At steady state, the oxygen concentration at any location within the well does not vary with time, so:

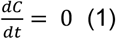

Furthermore, for concentration to remain constant, the rate of oxygen entering the well must balance the rate of oxygen consumption. The side wall of the well is assumed to be impermeable to oxygen. The sole location of entry for oxygen into the well is through the top surface (air-liquid boundary). This boundary is assumed to be constant concentration. The sole location of oxygen consumption is through the bottom surface, which models a cellular monolayer, and the consumption rate is assumed to be constant. Lastly, the diameter of the well is assumed to be constant along the depth of the well. Therefore, we may write the flux balance equation:

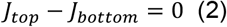

where *J*_*top*_ is the oxygen flux into the well from the top surface of the medium, and *J*_*bottom*_ is the oxygen consumption flux at the well bottom. Furthermore, because of the cylindrical symmetry, O_2_ concentration is dependent only on z, and the O_2_ concentration gradient and flux are constant throughout the volume. So *J*_*top*_ is given by Fick’s First Law as:

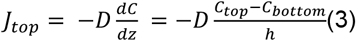

At the same time, *J*_*top*_ must also be equal to the cellular oxygen consumption rate (*OCR)* per unit area at the well bottom *J*_*bottom*_:

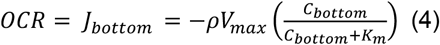

Substituting (3) and (4) into (2) yields

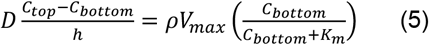

Equation (5) can be solved for pericellular oxygen C_bottom_ using the quadratic formula and keeping the positive root.

Rearranging (5) gives:

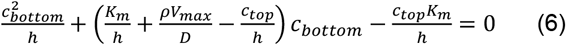

The dimensionless version of equation (6) is:

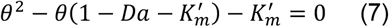

where,

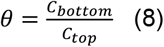

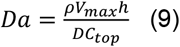

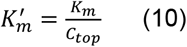

So

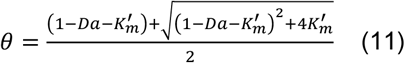

### Damkohler Number *Da* and Oxygen supply and consumption - Extreme cases when *Da* << 1 or *Da* >> 1

Note: for cardiomyocytes, *K*_*m*_*’* = *K*_*m*_/*c*_*to*p_ ∼ 5uM/179uM = 0.028 <<1.

(1) For *Da* << 1:

It can be shown from equation (11) that:

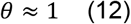

i.e.

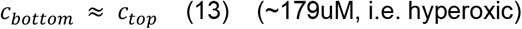

(2) For *Da* >> 1:

Equation (11) can be written as:

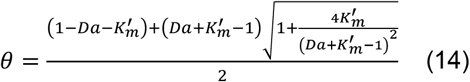

Using 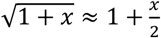, for *x* ≪ 1, eq. (14) is approximated as:

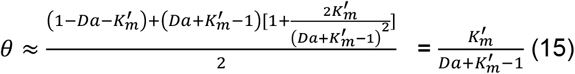

given *Da* >>1, so

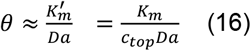

i.e.

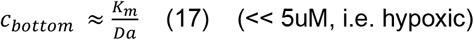

## REFERENCES

1. Ahmed, S. M., R. V. Shivnaraine and J. C. Wu (2023). “FDA Modernization Act 2.0 Paves the Way to Computational Biology and Clinical Trials in a Dish.” Circulation 148(4): 309–311.

2. Al-Ani, A., D. Toms, D. Kondro, J. Thundathil, Y. Yu and M. Ungrin (2018). “Oxygenation in cell culture: Critical parameters for reproducibility are routinely not reported.” PLoS One 13(10): e0204269.

3. Bagshaw, O. R. M., M. De Lange, S. Renda, A. J. F. Valente and J. A. Stuart (2019). “Hypoxio: a simple solution to preventing pericellular hypoxia in cell monolayers growing at physiological oxygen levels.” Cytotechnology 71(4): 873–879.

4. Barker, R. A., M. Carpenter, C. H. M. Jamieson, C. E. Murry, G. Pellegrini, R. C. Rao and J. Song (2022). “Lessons learnt, and still to learn, in first in human stem cell trials.” Stem Cell Reports.

5. Casey, T. M. and P. G. Arthur (2000). “Hibernation in noncontracting mammalian cardiomyocytes.” Circulation 102(25): 3124–3129.

6. Desa, D. E., T. Qian and M. C. Skala (2023). “Label-free optical imaging and sensing for quality control of stem cell manufacturing.” Curr Opin Biomed Eng 25.

7. Gerencser, A. A., A. Neilson, S. W. Choi, U. Edman, N. Yadava, R. J. Oh, D. A. Ferrick, D. G. Nicholls and M. D. Brand (2009). “Quantitative microplate-based respirometry with correction for oxygen diffusion.” Anal Chem 81(16): 6868–6878.

8. Grün, C., J. Pfeifer, G. Liebsch and E. Gottwald (2023). “O(2)-sensitive microcavity arrays: A new platform for oxygen measurements in 3D cell cultures.” Front Bioeng Biotechnol 11: 1111316.

9. Han, J. L., Y. W. Heinson, C. J. Chua, W. Liu and E. Entcheva (2023). “CRISPRi gene modulation and all-optical electrophysiology in post-differentiated human iPSC-cardiomyocytes.” Communications Biology 6(1): 1236.

10. Heinson, Y. W., J. L. Han and E. Entcheva (2023). “OptoDyCE-plate as an affordable high throughput imager for all optical cardiac electrophysiology.” Journal of Molecular and Cellular Cardiology Plus 6: 100054.

11. Jomezadeh Kheibary, N., J. Abolfazli Esfahani and S. A. Mousavi Shaegh (2020). “Analysis of oxygen transport in microfluidic bioreactors for cell culture and organ-on-chip applications.” Engineering Reports 2(1): e12062.

12. Keeley, T. P. and G. E. Mann (2019). “Defining Physiological Normoxia for Improved Translation of Cell Physiology to Animal Models and Humans.” Physiol Rev 99(1): 161–234.

13. Klimas, A., G. Ortiz, S. C. Boggess, E. W. Miller and E. Entcheva (2020). “Multimodal on-axis platform for all-optical electrophysiology with near-infrared probes in human stem-cell-derived cardiomyocytes.” Prog Biophys Mol Biol 154: 62–70.

14. Kohl, P. and R. G. Gourdie (2014). “Fibroblast-myocyte electrotonic coupling: Does it occur in native cardiac tissue?” J Mol Cell Cardiol.

15. Kostecki, G. M., Y. Shi, C. S. Chen, D. H. Reich, E. Entcheva and L. Tung (2021). “Optogenetic current in myofibroblasts acutely alters electrophysiology and conduction of co-cultured cardiomyocytes.” Sci Rep 11(1): 4430.

16. Li, W., J. L. Han and E. Entcheva (2022). “Protein and mRNA Quantification in Small Samples of Human-Induced Pluripotent Stem Cell-Derived Cardiomyocytes in 96-Well Microplates.” Methods Mol Biol 2485: 15–37.

17. Li, W., D. McLeod, J. T. Ketzenberger, G. Kowalik, R. Russo, Z. Li, M. W. Kay and E. Entcheva (2023). “High-throughput optical sensing of peri-cellular oxygen in cardiac cells: system characterization, calibration, and testing.” Frontiers in Bioengineering and Biotechnology 11.

18. Liu, W., J. L. Han, J. Tomek, G. Bub and E. Entcheva (2023). “Simultaneous Widefield Voltage and Dye-Free Optical Mapping Quantifies Electromechanical Waves in Human Induced Pluripotent Stem Cell-Derived Cardiomyocytes.” ACS Photonics 10(4): 1070–1083.

19. Magliaro, C., G. Mattei, F. Iacoangeli, A. Corti, V. Piemonte and A. Ahluwalia (2019). “Oxygen Consumption Characteristics in 3D Constructs Depend on Cell Density.” Front Bioeng Biotechnol 7: 251.

20. Marchiano, S., K. Nakamura, H. Reinecke, L. Neidig, M. Lai, S. Kadota, F. Perbellini, X. Yang, J. M. Klaiman, L. P. Blakely, E. Karbassi, P. A. Fields, A. M. Fenix, K. M. Beussman, A. Jayabalu, F. A. Kalucki, J. C. Potter, A. Futakuchi-Tsuchida, G. J. Weber, S. Dupras, H. Tsuchida, L. Pabon, L. Wang, B. C. Knollmann, S. Kattman, R. S. Thies, N. Sniadecki, W. R. MacLellan, A. Bertero and C. E. Murry (2023). “Gene editing to prevent ventricular arrhythmias associated with cardiomyocyte cell therapy.” Cell Stem Cell 30(4): 396-414.e399.

21. Mehta, G., K. Mehta, D. Sud, J. W. Song, T. Bersano-Begey, N. Futai, Y. S. Heo, M. A. Mycek, J. J. Linderman and S. Takayama (2007). “Quantitative measurement and control of oxygen levels in microfluidic poly(dimethylsiloxane) bioreactors during cell culture.” Biomed Microdevices 9(2): 123–134.

22. Nit, K., M. Tyszka-Czochara and S. Bobis-Wozowicz (2021). “Oxygen as a Master Regulator of Human Pluripotent Stem Cell Function and Metabolism.” J Pers Med 11(9).

23. Peniche Silva, C. J., G. Liebsch, R. J. Meier, M. S. Gutbrod, E. R. Balmayor and M. van Griensven (2020). “A New Non-invasive Technique for Measuring 3D-Oxygen Gradients in Wells During Mammalian Cell Culture.” Front Bioeng Biotechnol 8: 595.

24. Place, T. L., F. E. Domann and A. J. Case (2017). “Limitations of oxygen delivery to cells in culture: An underappreciated problem in basic and translational research.” Free Radical Biology and Medicine 113: 311–322.

25. Rog-Zielinska, E. A., R. A. Norris, P. Kohl and R. Markwald (2016). “The Living Scar--Cardiac Fibroblasts and the Injured Heart.” Trends Mol Med 22(2): 99–114.

26. Rolletschek, H. and G. Liebsch (2017). “A Method for Imaging Oxygen Distribution and Respiration at a Microscopic Level of Resolution.” Methods Mol Biol 1670: 31–38.

27. Sekine, K., Y. Kagawa, E. Maeyama, H. Ota, Y. Haraguchi, K. Matsuura and T. Shimizu (2014). “Oxygen consumption of human heart cells in monolayer culture.” Biochem Biophys Res Commun 452(3): 834–839.

28. Song, Q., A. Alvarez-Laviada, S. E. Schrup, B. Reilly-O’Donnell, E. Entcheva and J. Gorelik (2023). “Opto-SICM framework combines optogenetics with scanning ion conductance microscopy for probing cell-to-cell contacts.” Commun Biol 6(1): 1131.

29. Tan, J., S. Virtue, D. M. Norris, O. J. Conway, M. Yang, G. Bidault, C. Gribben, F. Lugtu, I. Kamzolas, J. R. Krycer, R. J. Mills, L. Liang, C. Pereira, M. Dale, A. S. Shun-Shion, H. J. M. Baird, J. A. Horscroft, A. P. Sowton, M. Ma, S. Carobbio, E. Petsalaki, A. J. Murray, D. C. Gershlick, J. A. Nathan, J. E. Hudson, L. Vallier, K. H. Fisher-Wellman, C. Frezza, A. Vidal-Puig and D. J. Fazakerley (2024). “Limited oxygen in standard cell culture alters metabolism and function of differentiated cells.” The EMBO Journal: 1-39-39.

30. Truskey, G., F. Yuan and D. Katz (2009). Transport Phenomena in Biological Systems, 2e, Pearson, ch 7, pp 496–514.

31. Tschiersch, H., G. Liebsch, L. Borisjuk, A. Stangelmayer and H. Rolletschek (2012). “An imaging method for oxygen distribution, respiration and photosynthesis at a microscopic level of resolution.” New Phytol 196(3): 926–936.

32. Veldhuizen, J., R. Chavan, B. Moghadas, J. G. Park, V. D. Kodibagkar, R. Q. Migrino and M. Nikkhah (2022). “Cardiac ischemia on-a-chip to investigate cellular and molecular response of myocardial tissue under hypoxia.” Biomaterials 281: 121336.

33. Wagner, B. A., S. Venkataraman and G. R. Buettner (2011). “The rate of oxygen utilization by cells.” Free Radic Biol Med 51(3): 700–712.

34. Ward, M. C. and Y. Gilad (2019). “A generally conserved response to hypoxia in iPSC-derived cardiomyocytes from humans and chimpanzees.” Elife 8.

35. Wei, L., W. Li, E. Entcheva and Z. Li (2020). “Microfluidics-enabled 96-well perfusion system for high-throughput tissue engineering and long-term all-optical electrophysiology.” Lab Chip 20(21): 4031–4042.

36. Ye, L., L. Qiu, B. Feng, C. Jiang, Y. Huang, H. Zhang, H. Zhang, H. Hong and J. Liu (2020). “Role of Blood Oxygen Saturation During Post-Natal Human Cardiomyocyte Cell Cycle Activities.” JACC: Basic to Translational Science 5(5): 447–460.

